# On the post-glacial spread of human commensal *Arabidopsis thaliana*: journey to the east

**DOI:** 10.1101/411108

**Authors:** Che-Wei Hsu, Cheng-Yu Lo, Cheng-Ruei Lee

## Abstract

With the availability of more sequenced genomes, our understanding of the evolution and demographic history of the model plant *Arabidopsis thaliana* is rapidly expanding. Here we compile previously published data to investigate global patterns of genetic variation. While the Southeast African accessions were reported to be the most divergent among worldwide populations, we found accessions from Yunnan, China to be genetically close to the sub-Saharan accessions. Our further investigation of worldwide chloroplast genomes identified several deeply diverged haplogroups existing only in Eurasia, and the African populations have lower variation in many haplogroups they shared with the Eurasian populations. Bayesian inferences of chloroplast demography showed that representative haplogroups of Africa exhibited long-term stable population size, suggesting recent selective sweep or bottleneck is not able to explain the lower chloroplast variation in Africa. Taken together, these patterns cannot be easily explained by a single out-of-Africa event. Several Eurasian chloroplast haplogroups had rapid population growth since 10 kya, presumably reflecting the recent expansion of the weedy non-relicts across Eurasia. Our demographic analysis on a chromosomal region un-affected by relict introgression also suggested the European, Central Asian, and Chinese Yangtze populations diverged no earlier than 15 kya, in contrast to previous estimates of 45 kya inferred from whole genome that likely contains relict admixture. The most recent expansion is observed in the Yangtze population of China less than 2000 years ago. Similar to Iberia, the western end of non-relict expansion reported in our previous study, in this eastern end of Eurasia we find clear traces of gene flow between the Yangtze non-relicts and the Yunnan relicts. Genes under strong selection and previously suggested to contribute to adaptation in the Yangtze valley are enriched for traces of relict introgression, especially those related with biotic and immune responses. The results suggest the ability of non-relicts to obtain locally adaptive alleles through admixture with relicts is an important factor contributing to the rapid expansion across the environmental gradients spanning the eastern to the western coast of Eurasia.

## Introduction

*Arabidopsis thaliana* is not only a model species in plant molecular biology, but also increasingly used to address major questions in ecology and evolution. The evolutionary history of this model plant is frequently revisited as more geographically diverse samples are collected and sequenced. The first continental-scale study of species-wide demography is published in 2016, where one globally distributed human commensal “non-relict” group as well as several “relict” groups located in relatively un-disturbed habitats were identified^1^. The follow-up study showed that the non-relicts originated recently near the Balkans and spread along the east-west axis of Eurasia, wiping out continental-wide relict populations while incorporating locally adaptive alleles from them^2^. As to the new world, the North American population arrived at around 1600 AD^3^, and most of them likely came from the region near southeastern England and northwestern Germany, carrying a charismatic inversion in chromosome 4^4^.

Durvasula *et al.*^5^ investigated African *A. thaliana* and suggested an “out of Africa” demographic model, given the highest genomic variation and numbers of private alleles observed in African accessions. In this model, *A. thaliana* originated in Africa and diverged into three populations at ca. 90 kya: The Moroccan, Levantine and Southeast African groups. The migration of Moroccan population northwards to Iberia was illustrated as well as the Levantine migration wave westward into Europe and eastward to Central Asia at ca. 45 kya^6^. On the other hand, it remains unclear how *A. thaliana* first arrived Africa given all other *Arabidopsis* species were found in temperate Northern Hemisphere^7^, and the pattern that Africa contains most variation can be equally likely explained by a non-African origin of *A. thaliana* followed by non-relict expansion wiping out most Eurasian variation^2^.

Zou *et al.*^8^ studied *A. thaliana* accessions from China and showed that the population in Yangtze River Basin arrived relatively recently. They also showed that genes associated with immune response as well as flowering time were significantly enriched in the list of selected genes in the Yangtze population. For flowering time, genetic mapping identified a candidate gene in chromosome 2, containing the *SVP* gene (AT2G22540) with a loss-of-function mutation accelerating flowering. It remains unclear what constitutes the source of adaptive allele in the Yangtze population – the sources of adaptation may be novel mutation, standing variation, or as we have shown for the Iberian non-relicts, introgression from locally adaptive relicts^2^.

Here we compile global data and re-investigate the evolutionary history of *A. thaliana* with two specific aims. (1) We used the maternally inherited chloroplast genomes to study the species history from a different perspective and investigate the out-of-Africa hypothesis in the context of global samples. (2) We wish to clear up the evolutionary history and timing of non-relict expansion across Eurasia, especially whether the Chinese Yangtze population represents the eastern end of non-relict expansion and whether adaptive introgression also happened there.

## Results

### Genetic variation in nuclear genomes

To investigate the global patterns of *Arabidopsis thaliana* genomic variation, we compiled data from the 1,001 genomes project^1^, the African accessions^5^, and the Chinese accessions^8^. Phylogenetic tree using *Arabidopsis lyrata* as the outgroup^7^ confirmed the previous observation that the Tanzanian and South African accessions (hereafter the “TZSA” clade) are most divergent to all others (Fig. 1a, Supplementary Fig. 1). Interestingly, two accessions from Yunnan, China are also genetically close to the TZSA clade, inconsistent with the single out-of-Africa event suggested previously^5,6^.

**Fig. 1.**
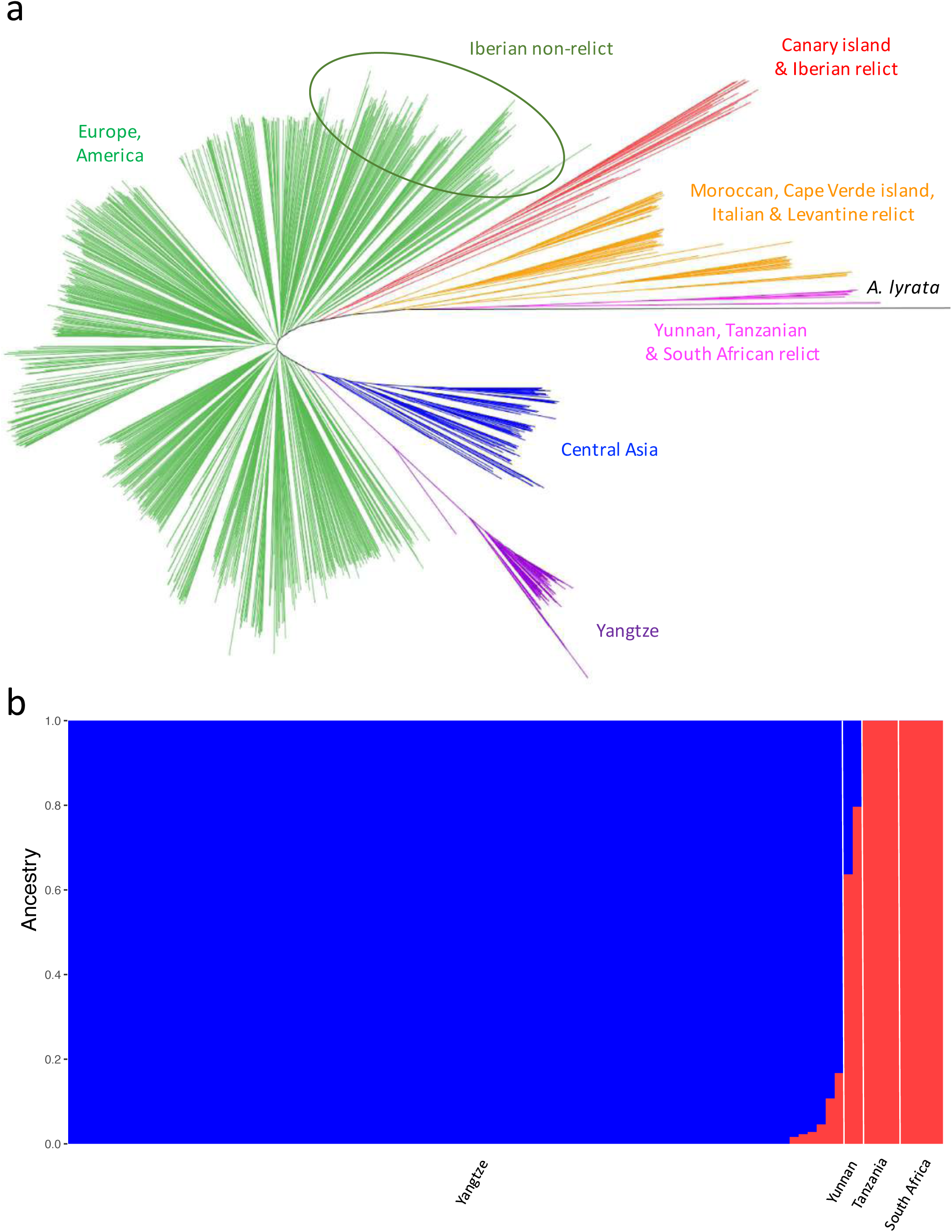
Differentiation of *Arabidopsis thaliana* nuclear genomes. (a) Neighbor-Joining tree. (b) K=2 ADMIXTURE result.

Of the two Yunnan accessions, one (SRR2204703) has heterozygosity typical of self-fertilizing *A. thaliana*, and the other (SRR2204316) has very high heterozygosity (Supplementary Fig. 2). While the higher heterozygosity may result from recent outcrossing events or DNA contamination, both samples have very low chloroplast heterozygosity as all other accessions (< 0.001), making DNA contamination less likely. Since the number of heterozygous sites in an individual reflects the number of SNPs between its two parents, we suspected the high heterozygosity of SRR2204316 might result from the cross between two genetically divergent groups, similar to a recent study in ancient humans^9^.

The ADMIXTURE^10^ K = 2 result supports this idea (Fig. 1b). While the Chinese Yangtze population is highly similar to typical Eurasian non-relicts and the Yunnan accessions are close to the TZSA group (Fig. 1a), admixture exists (Fig. 1b). We further investigated this with ABBA-BABA tests (Table 1). We first used non-relicts from Western Europe, which in theory had no gene flow with any of the Tanzanian/South African/Yunnan relict population, as a reference group to test whether the Yunnan accessions had gene flow from non-relicts. The sign of gene flow is highly significant for the highly heterozygous Yunnan accession (*P* = 5.17E-25, Table 1 Test A) but not the other (*P* = 0.539, Table 1 Test A). Using the relatively un-admixed Yunnan accession as a reference, the Chinese Yangtze non-relicts showed strong signs of gene flow from the Yunnan relicts (Table 1 Test B). Finally, since both the highly heterozygous Yunnan accession and the Chinese Yangtze population showed signs of admixture, the tests involving both groups are highly significant (Table 1 Test C). Therefore, similar to Iberia, the far-eastern end of Eurasia is also affected by the rapid expansion of non-relict population, with introgression from local relicts along the way.

**Table 1.** Results from the ABBA-BABA test in the form of (((P1,P2),P3),O) where O is the outgroup *Arabidopsis lyrata* ^a^

| Test | P1 | P2 | P3 | *D* statistic | Z score | *P* value |
| --- | --- | --- | --- | --- | --- | --- |
| A | TZSA | YU-admix | EU | 0.172 | 10.330 | 5.17E-25 |
| A | TZSA | YU-pure | EU | 0.011 | 0.614 | 0.539 |
| B | EU | YA | YU-pure | 0.081 | 3.967 | 7.27E-05 |
| B | TZSA | YU-pure | YA | 0.063 | 3.023 | 0.003 |
| C | EU | YA | YU-admix | 0.256 | 14.562 | 4.92E-48 |
| C | TZSA | YU-admix | YA | 0.365 | 21.756 | 6.05E-105 |

a. EU: Western European non-relicts. TZSA: Tanzanian and South African relicts. YA: Chinese Yangtze River Basin non-relicts. YU-admix: The more-admixed Yunnan relict with high heterozygosity. YU-pure: The less-admixed Yunnan relict.

### Genetic variation in chloroplast genomes

Our results from the nulcear genome suggest a more complex demography than a single out-of-Africa event^5,6^. To better understand this, we investigated the chloroplast genomes, dated with 11 outgroup species^7,11^. Principal component analysis (PCA) of chloroplast variation within *A. thaliana* identified several genetic groups (Fig. 2a, Supplementary Fig. 3), which is consistent with the phylogenetic tree (Supplementary Fig. 4). After collapsing branches with low approximate likelihood ratio support (Supplementary Fig. 5), we found that the *A. thaliana* chloroplast tree exhibits a basal polytomy (Fig. 2b), with seven major haplogroups branched off at roughly the same time: groups 1, 2, 8, 9, 10, and two monophyletic clades: 3 + 4 and 5 + 6 + 7. While studies based on the nuclear genomes showed that the African accessions contain higher polymorphism and are highly diverged from Eurasian populations^5^, we did not observe such pattern in chloroplast. Instead, the African accessions represent only a small subset of chloroplast variation (Groups 2, 3, and 7, Fig. 2b), suggesting that the chloroplast phylogeny captures more ancient demographic history than the divergence between African and European populations. Indeed, molecular dating with BEAST^12^ confirmed that the major chloroplast haplogroups diverged at ca. 227 kya (95% highest posterior density 121-340 kya, Supplementary Fig. 6,7). This considerably predates the inferred divergence time between African and Eurasian nuclear genomes (90-120 kya)^5,6^.

**Fig. 2.**
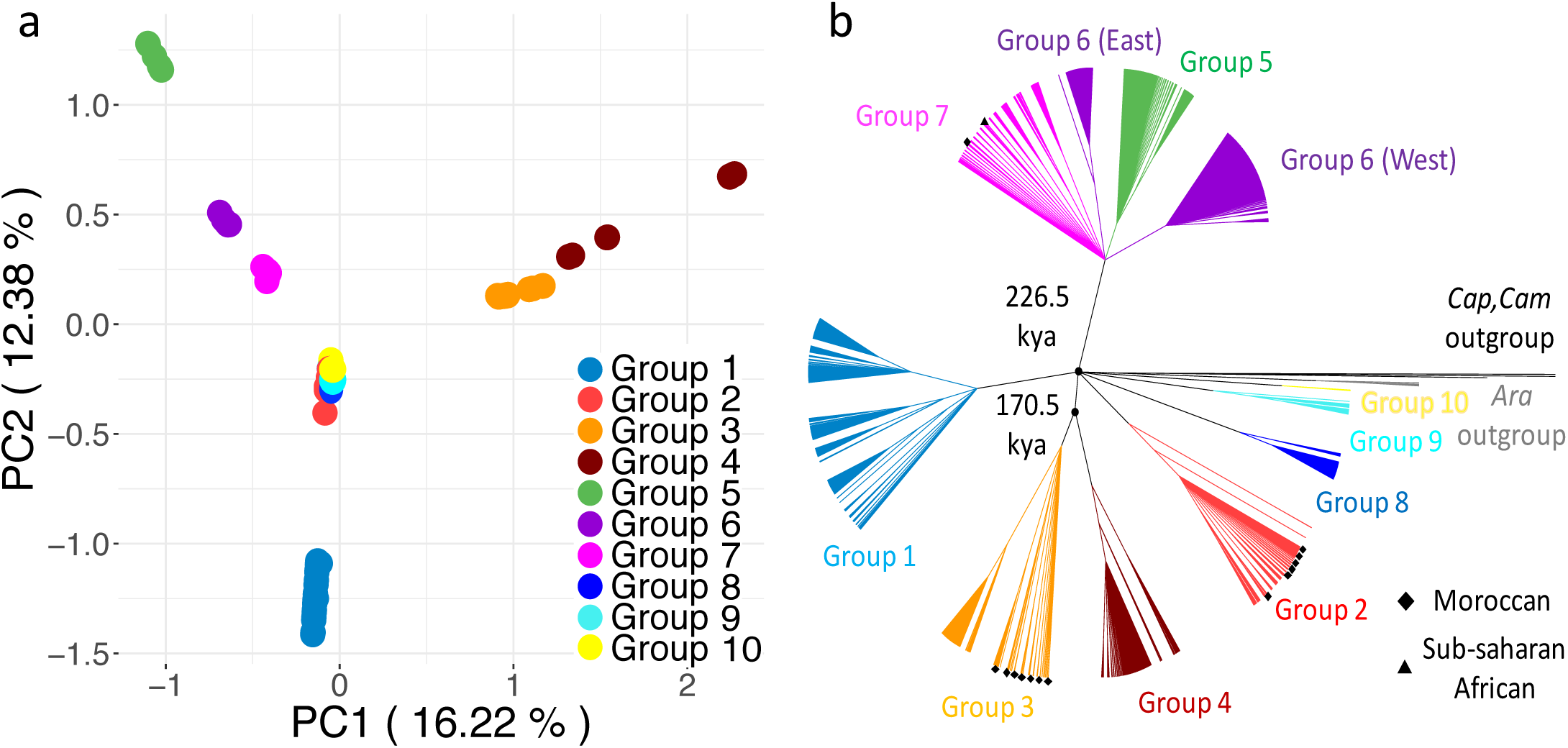
Differentiation of *Arabidopsis thaliana* chloroplast genomes. (a) Principal component analysis. (b) Maximum likelihood cladogram, where branches with low aLRT support were collapsed. Group 6 (East) consists of samples from Yangtze River Basin, China and Kashmir, India. Group 6 (West) consists of samples in Eurasia.

The spatial distribution of chloroplast haplogroups is uneven, with several highly diverged haplogroups existing only in Eurasia, and the African population containing only group 2, 3, and 7 (Fig. 3a). While Morocco has highest nuclear genomic variation^5^, we observed this only for group 3, where the variation decreases from Morocco northwards (Fig. 3c), suggesting its Moroccan origin and later northward migration. Group 2 is only confined in Iberia and Morocco, with the former having higher variation (Fig. 3b). The monophyletic clade containing group 5, 6 and 7, on the other hand, has a global distribution with highest variation in Europe, especially the Balkan Peninsula (Fig. 3e). Therefore, even among the three haplogroups Africa shares with Eurasia, only one of them has African population containing higher variation. In summary, our observation is consistent with both hypotheses of (1) African origin of *A. thaliana* and complete lineage sorting between the African and the Eurasian accessions or (2) a Eurasian origin and dispersal into different regions (southwestern Europe and northwestern Africa, Balkan and Levant, and south Asia), after which most Eurasian nuclear genomic variation was wiped out by the rapidly expanding non-relicpts.

**Fig. 3.**
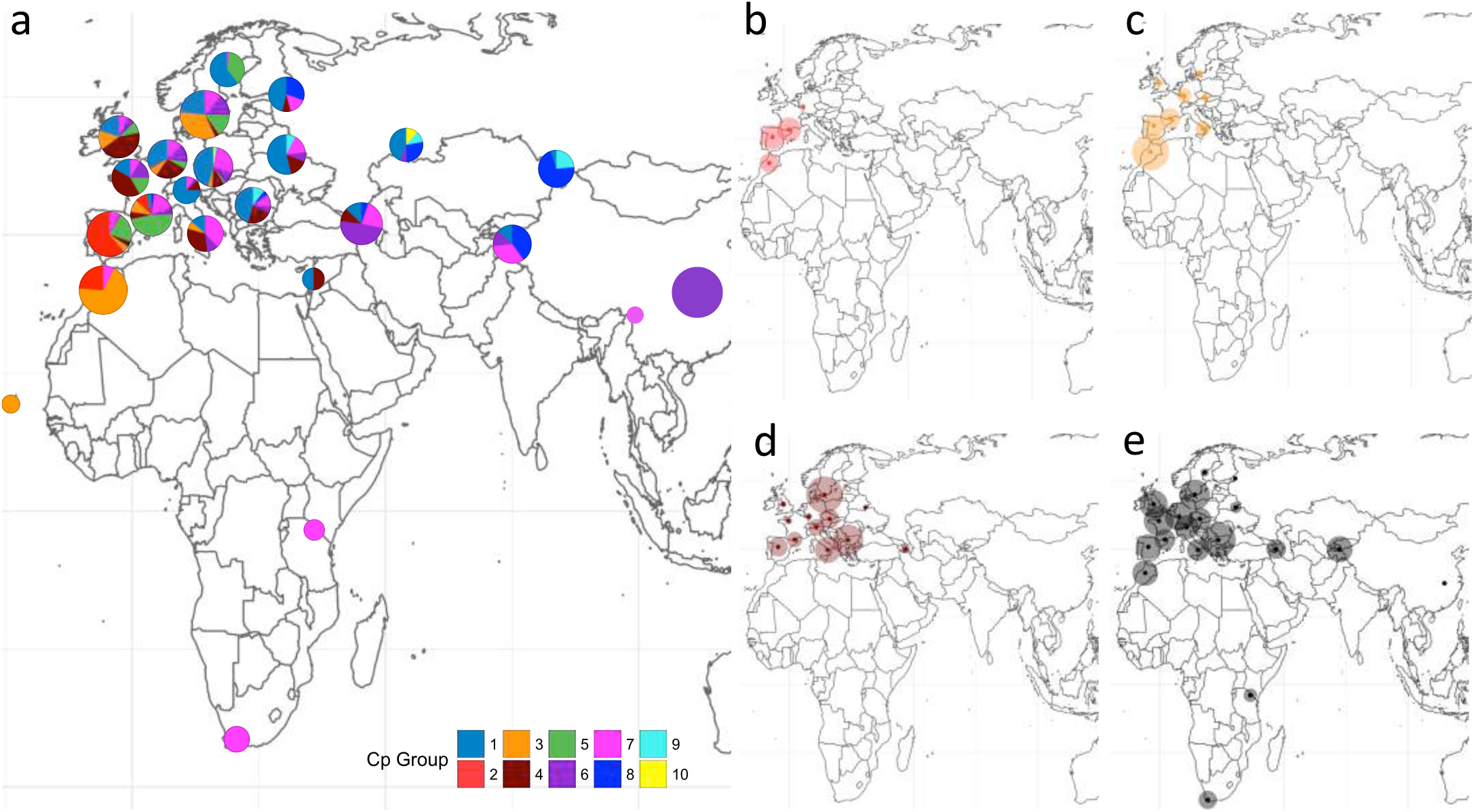
Geographical distribution and spatial genetic variation of chloroplast haplogroups. (a) Diversity map of chloroplast haplogroups. Pie charts show the proportion of each group, and chart size is proportional to sample size. (b), (c), (d), (e) are polymorphism maps correspond to group 2, 3, 4, 5+6+7 respectively. The diameter of each circle is proportional to mean pair-wise genetic distance of each geographical region.

### Demography of chloroplast genomes

We further used Extended Bayesian Skyline Plots^13^ to investigate chloroplast demographic histories. Haplogroups typical of the Iberian and Moroccan region (group 2, 3, and 4) showed long-term stable population size (Fig. 4). The Eurasian haplogroups 1 and 5 had population size increase since 20 kya, consistent with the post-glacial expansion with the retreat of ice sheet. Interestingly, for the globally distributed group 7, the central Asian group 8, the European group 6W, and the Chinese Yangtze group 6E, all had rapid population size increase since 10 kya, a time point close to the previously inferred rapid expansion of weedy non-relicts^2,14^. In addition, haplogroups 6 and 7 had highest genetic variation near the Balkan Peninsula (Supplementary Fig. 8c,d), corresponding to the inferred origin of non-relict expansion^2^.

**Fig. 4.**
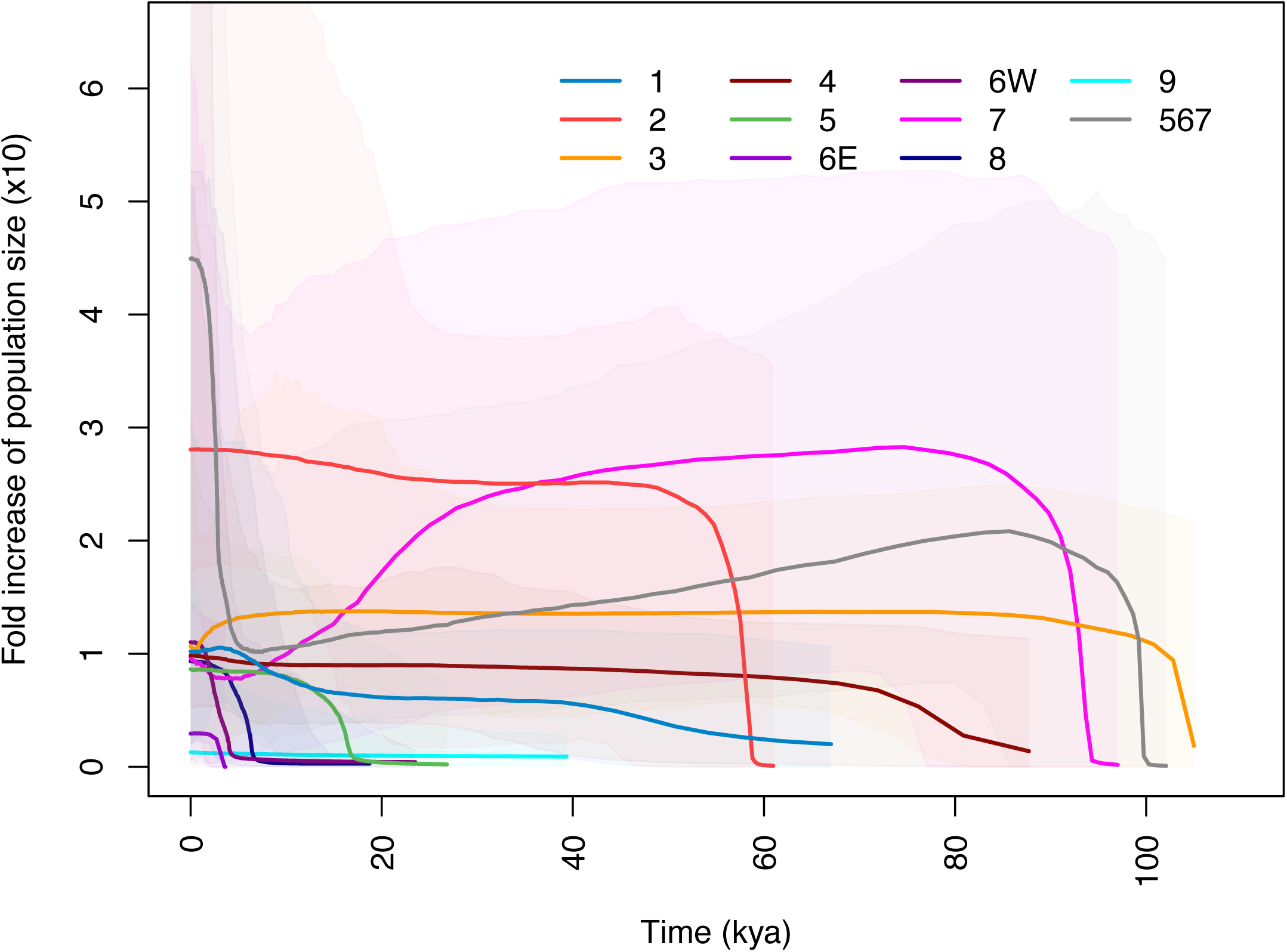
Variation of population size over time inferred from chloroplast polymorphism. Extended Bayesian Skyline is plotted for each chloroplast haplogroup except group 10, which has small sample size.

For haplogroups shared by Morocco and Europe (groups 2, 3, and 7), we further investigated their demographic histories separately. While the Morocco population of all three groups still exhibited long-term stable population size (Supplementary Fig. 9), the European group 2 had population size increase since 20 kya (the first post-glacial expansion), and the European group 7 had size increase since 10 kya (the non-relict expansion). Taken together, compared to Europe, the Moroccan population was less influenced by either episode of demographic change, especially the second expansion wave wiping out most nuclear genomic variation across Eurasia^2^. While the fact that Morocco possesses most nuclear genomic variation could be interpreted as an African origin of *Arabidopsis thaliana*^5,6^, the complex demographic history we showed is an equally likely explanation.

### The eastern end of non-relict expansion

While all current and previous^2,14^ estimates suggested the non-relicts expanded around 10 kya and almost all Eurasian *A. thaliana* are descendants of this population, some studies estimated the population divergence time between the European, Central Asian, and Chinese non-relict populations to be around 45 kya^5,6,8^. While these studies performed the multiple sequentially Markovian coalescent (MSMC) estimates^15^ on the whole genome, we wish to note there are clear evidences that non-relict populations across Eurasia had introgression from distinct and highly diverged local relicts (Table 1 and ref. 2). Using the whole genome, in some genomic regions one would be comparing between relicts and non-relicts or two highly diverged relict groups (e.g. between Tanzania and Morocco), thereby overestimating the true divergence time between the European, Central Asian, and Chinese non-relict populations. We therefore focus on a unique chromosomal translocation, where the non-relicts have a charismatic derived haplotype^2,16^ (Supplementary Fig. 10). Since the two structural variants of the translocation cannot recombine effectively, genetic variation within the derived haplotype reflects demographic history of non-relicts^2^. Using only this 750 kb region, we first performed the same set of MSMC comparisons as Durvasula *et al.*^5^ and successfully re-created similar patterns (Supplementary Fig. 11), demonstrating this region alone contains enough information to trace the demographic history. Based on the derived haplotypes in this region, the European, Central Asian, and Chinese Yangtze non-relict populations diverged between 5 to 15 kya (Fig. 5), consistent with the time of rapid population expansion in several chloroplast haplogroups (10 kya, Fig. 4) as well as the inferred timing of non-relict expansion^2,14^.

**Fig. 5.**
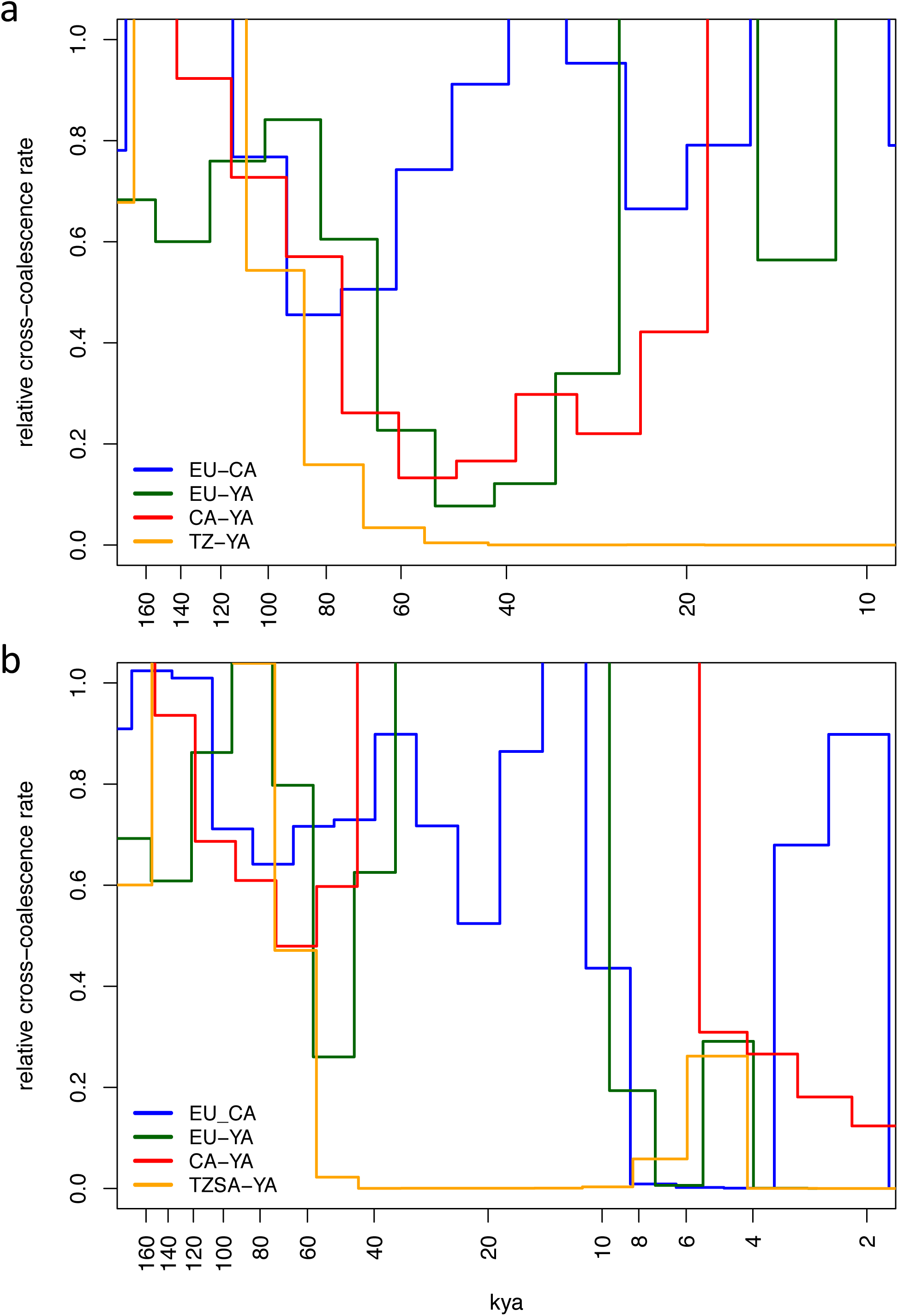
Timing of population splits inferred from chromosome 1 translocation. Relative cross coalescence rate (CCR) between populations is shown. EU: Western Europe, CA: Central Asia, YA: Yangtze, TZ: Tanzania, TZSA: Tanzania and South Africa. Decrease of CCR from 1.0 indicates population split, steep slope of CCR from 1.0 to 0.0 indicates drastic and complete isolation while mild one indicates slow and progressive isolation. (a) 4 haplotypes MSMC that has better estimation of older splits. (b) 8 haplotypes MSMC that has better estimation of more recent splits.

For non-relicts, the most notable recent expansion happened in the Chinese Yangtze population (Fig. 4). Assuming the North American population had a common ancestor around 1600 AD^3^, using simple genetic distance and assuming the same mutation rate, we estimated the Chinese Yangtze population having a common ancestor at 568 AD (estimated from chloroplast) or 823 AD (estimated from the chromosomal translocation region in chromosome 1^2^). The time point is consistent with *A. thaliana* entering China through central Asia with human activities, with the Silk Road being one possibility.

Given that the Yangtze population clearly had introgression from the Yunnan relicts (Fig. 1b and Table 1), we further investigated whether introgression contributed to local adaptation of this population. We calculated the 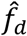 statistic^17^ for 50-kb windows across the genome, from which gene flow between Yangtze population and Yunnan relicts was inferred (Supplementary Table 1). Then we compared the results with genes under strong selection in Yangtze population identified by Zou *et al.*^8^ (Supplementary Table 1,2). Windows with these selected genes were found enriched in the top 5% tail of 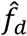 distribution (Fisher’s exact test, odds ratio = 2.049, *P* = 0.007). Genes both under strong selection and with evidences of relict origin were overrepresented for gene ontology (GO) terms associated with biotic interaction, immune response, and programmed cell death (Supplementary Table 2), while strongly selected genes without strong traces of introgression (presumably representing novel mutations or standing variations within the invading ancestral Yangtze non-relicts) have no enrichment of any GO term. Taken together, much similar to the western end of Eurasia^2^, our results suggested the ability of the expanding non-relict population to colonize the eastern end of Eurasia (the Yangtze River Basin) was also greatly facilitated by introgressions from local relicts.

Interestingly, in whole-genome phylogenetic tree the Yangtze population has long branches relative to other non-relict populations (Fig. 1a). To test whether this is caused by introgression from a highly diverged group (the Yunnan relicts) or natural selection accelerating the fixation of novel mutations in some genomic regions, we excluded the top 20% windows with highest introgression (Supplementary Fig. 12a), any window containing positively selected genes (Supplementary Fig. 12b), or both (Fig. 12c). These trees remain similar to the whole-genome tree where the Yangtze population still has long branch length. It is likely that the Yangtze population exhibits higher mutation rate or more rapid life cycle resulting in more than one generation per year, and both hypotheses need to be formally tested. If so, time to the most recent common ancestor of Yangtze accessions would be more recent than our estimation.

## Discussion

### On ancient population structure

Combining currently available data from genome resequencing projects of *Arabidopsis thaliana*, here we revisit demographic history of *A. thaliana* from the global perspective. The “out of Africa” hypothesis states that the African population first separated into the Moroccan, Levantine, and Southeast African groups at ca. 90 kya followed by a migration event from Levant into Eurasia^5,6^. However, we observed that the Chinese Yunnan accessions are genetically closer to the Tanzanian and South African group than to any other group, which suggests more than one “out of Africa” events if the ancestral population is originated from Africa. For chloroplast, we observed several highly diverged haplogroups existing only in Eurasia, and Africa contains merely a subset of overall chloroplast variation, which hints that the ancestral population may not originate from Africa. Together, these results suggest another demographic scenario that is as possible as the “out of Africa” model (Fig. 6): Like all other species in the *Arabidopsis* genus, ancient *Arabidopsis thaliana* originated in temperate Eurasia and separated into the Moroccan/Iberian, Levantine, and South/Southwest Asian groups at ca. 90 kya. Later the Moroccan/Iberian and Levantine group migrated northwards into Eurasia while the Asian group dispersed into Tanzania and South Africa. At around 10 kya, the weedy non-relict group expanded across Eurasia. On the other hand, we acknowledge while the existence of more ancient chloroplast variation in Eurasia might indicate a Eurasian origin of *A. thaliana*, it is also likely that the ancient variation once existed in Africa but was later lost due to strong bottleneck events or selective sweeps favoring a few chloroplast haplogroups. Chloroplast demography, however, shows long-term stable population size in Morocco for all haplogroups (Supplementary Fig. 9), thus not lending strong support to the Moroccan chloroplast bottleneck or sweep scenario.

**Fig. 6.**
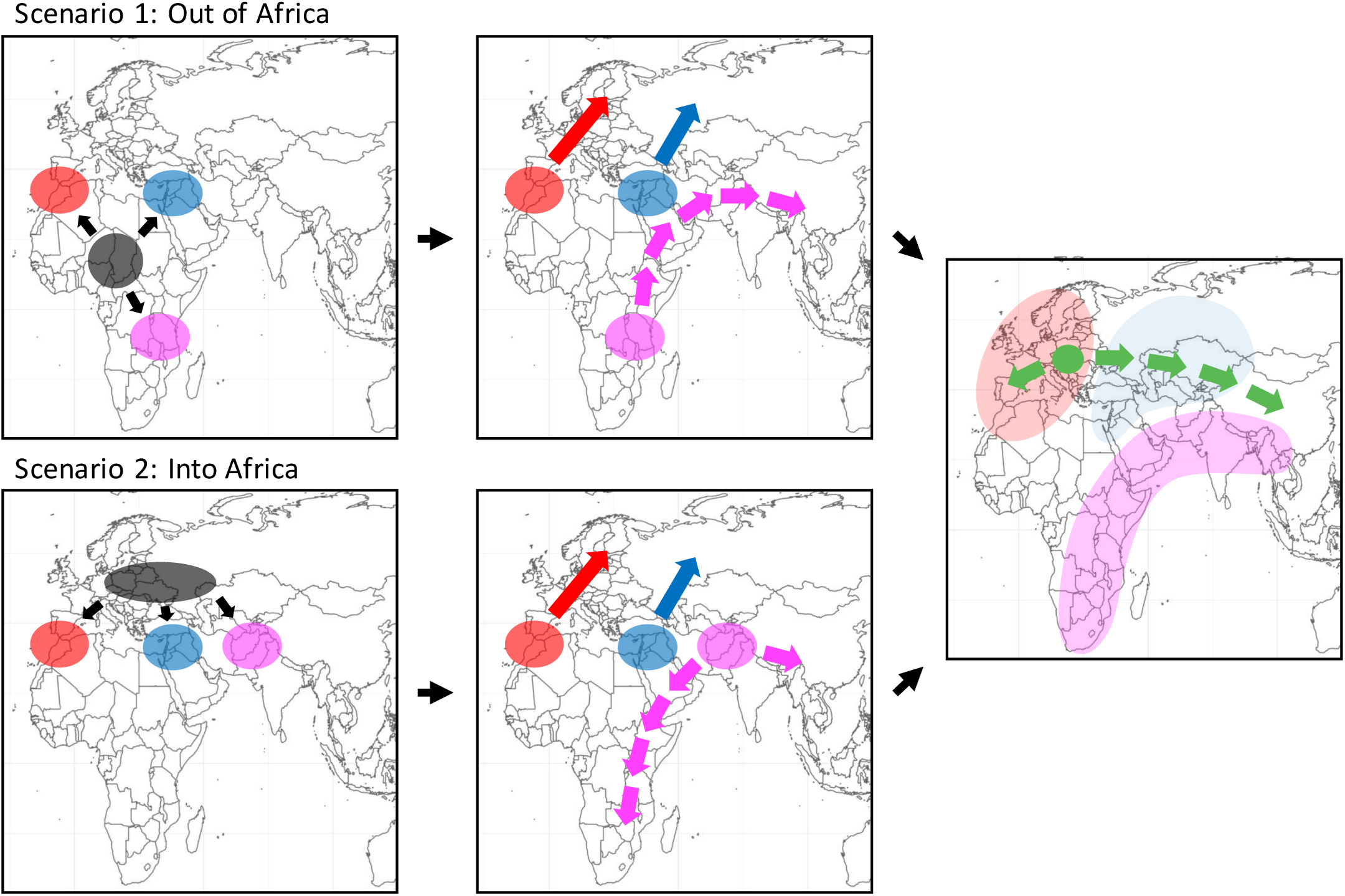
Two scenarios of demographic history consistent with the present-day pattern of spatial genetic variation in *Arabidopsis thaliana*. Scenario 1 represents the “Out of Africa” model: Ancestral population of *Arabidopsis thaliana* in Africa split into 3 populations at ca. 90 kya, the Moroccan, Levantine and Sub-saharan African. Later, Moroccan expanded into Europe through Iberia, Levantine dispersed into Central Asia and Europe while Sub-saharan African migrated into Yunnan possibly through Southwest and South Asia. Scenario 2 represents the “Into Africa” model: Ancestral population of *Arabidopsis thaliana* in Europe split into 3 populations, the Moroccan/Iberian, Levantine and a South/Southwest Asian population. Later, Moroccan/Iberian expanded northwards and eastwards into Europe, Levantine dispersed into Central Asia and Europe while the South/Southwest Asian migrated into Yunnan and Sub-Saharan Africa. Since 10 kya, the weedy non-relicts from Balkan and Eastern Europe spread rapidly westwards into Iberia and eastwards into Yangtze River Basin of China, wiping out genetic variation along the way while obtaining adaptive genes through gene flow between local relicts.

For both the “out of Africa” and the “into Africa” models, the most notable demographic turnover event is the recent replacement of many Eurasian relict populations by the weedy “non-relicts"^1,2^, which expanded along the east-west axis of Eurasia and left more relict genomic fragments in southern and northern Europe. Interestingly, this is supported by an independent study: Exposito-Alonso *et al.*^18^ found that the same alleles increasing survival under extreme drought are enriched in the relict accessions and are concentrated in northern and southern Europe, a pattern predicted by the east-west non-relit expansion. The drastic demographic turnover is also responsible for only few private variants being observed in each regional Eurasian population^5^, which is a logical outcome since most Eurasian genomes descended only recently from a single population^1,2^. Meanwhile, the African population remained relatively isolated from Eurasia and retained much of the ancestral nuclear-genome variation. Therefore, our results suggest that the current patterns of global nuclear-genome variation (Africa containing most variation) is a consequence of non-relicts wiping out most ancient genetic variation in Eurasia^2^, which is compatible with both scenarios about ancient *A. thaliana* population structure (Fig. 6).

While no relict accession was found in northern central Asia, this region is enriched for the ancestral haplotype of the chromosome 1 translocation (Supplementary Fig 10), and these ancestral haplotypes form a unique genetic group when comparing to worldwide ancestral haplotypes^2^. In the present study, several Eurasia-only chloroplast haplogroups (group 8, 9, 10) also exist in this region (Fig 3). We therefore suspect another unique relict group might have existed near this region, which later became very rare or extinct.

### On the recently established Chinese population

In the process of rapid expansion, populations in the expansion front constantly encounter novel environments, which may impede the speed and extent of expansion. Hence, how can a population rapidly spread across a wide geographical and environmental range, and what is the source of adaptation to these drastically different environments? We therefore focus on the origin and adaptation of Chinese Yangtze population for the second part of our investigation. We show that the Yangtze population originated no more than 2000 years ago and spread rapidly across the basin. Using the properly rooted phylogenetic tree, we showed that Yangtze population belongs to the non-relict group and are genetically the closest to Central Asian non-relicts (Fig. 1).

Zou *et al.*^8^ performed genome-wide scans for signal of selection in the Yangtze population. Here we also investigated this result in the context of introgression from Yunnan relicts. We found that selected genes are enriched with signs of relict introgression, and genes with both signs of selection and introgression are overrepresented for immune-related functions. On the other hand, selected genes without signs of introgression do not have any significant gene ontology enrichment. Our results therefore suggest, among the various aspects of adaptation to the novel Yangtze River Basin environment, the adaptation to immune-related biotic stress is associated with gene flow with local relicts, which might have co-existed with local pathogens for a long time. Interestingly, for the western end of non-relict expansion in Iberia, relict introgression likely contributed to the adaptation to abiotic factor, as highly introgressed genes in Iberian non-relicts are enriched for GO terms including root development and ion metal transmembrane activity^2^. In the end, how can the non-relicts, a population near the Balkans, occupy such broad environmental gradient spanning more than 10,000 km across Eurasia within 10,000 years? While the mal-adaptation to novel environments in the expansion front may impede non-relict spread, our results suggest non-relicts frequently assimilated the biological distinctiveness of locally adaptive relicts. Together with human’s long-term disturbance of native Eurasian vegetation and non-relicts’ association with anthropogenically disturbed habitats^1,2^, the environmental resistance to non-relict expansion appears futile in most of Eurasia.

## Materials and Methods

### Data source and SNP identification of nuclear genome

In this study, we obtained *Arabidopsis thaliana* data from the 1,135 worldwide genomes^1^, 73 African genomes^5^, 116 Chinese genomes^8^ and one *Arabidopsis lyrata* sample (SRR2040792)^7^. Reads were trimmed based on quality using SolexaQA^19^, and possible remaining adaptor sequences were removed with cutadapt^20^. Reads were mapped to the TAIR 10 reference genome using BWA 0.7.15^21^. Picard Tools (http://broadinstitute.github.io/picard) were used to mark duplicated read pairs, and the genotypes of each site in each accession (including non-variant sites and SNPs) were called following GATK 3.7 best practice^22^.

We further filtered the SNPs with QUAL < 100, QD < 20, call rate < 0.99, DP < 3 or > 2 standard deviations from genome-wide average depth and removed 2 Chinese accessions (SRR2204178, SRR2204343) with high missing rate, resulting in 5,915,870 SNPs and 1323 accessions.

### Alignment of *A. thaliana* population data with outgroups

In addition to the *Arabidopsis thaliana* reference chloroplast genome (NC000932), we obtained the outgroup chloroplast genomes from the genera *Arabidopsis, Capsella*, and *Camelina*^7,11^: *Arabidopsis lyrata* subsp. *petraea* (LT161948), *Arabidopsis lyrata* subsp. *lyrata* (LN877383), *Arabidopsis halleri* subsp. *halleri* (LN877382), *Arabidopsis carpatica* (LT161918), *Arabidopsis arenosa* subsp. *arenosa* (LT161904), *Arabidopsis nitida* (LT161970), *Arabidopsis pedemontana* (LN877384), *Arabidopsis cebennensis* (LN877381), *Capsella rubella* (LN877385), *Capsella bursa-pastoris* (NC_009270), and *Camelina sativa* (LN877386).

All twelve chloroplast genomes were annotated with Verdant^23^, and about 90 protein coding genes were identified in each sequence. We retained genes existing in all species and excluded those within the two inverted repeat regions, resulting in 67 orthologous genes with one-to-one relationship in all species. The 67 genes were separately aligned with MUSCLE 3.8.31^24^. Based on this alignment of *A. thaliana* reference chloroplast genome with outgroups species, we used custom R scripts to “paste” the Illumina-based *A. thaliana* accession data^1,5,8^ onto the among-species alignment. All following analyses were based on this concatenated dataset of 67 protein coding genes.

To remove possible confounding effect from heteroplasmy, we excluded accessions with heterozygous SNPs > 0.5% of all SNPs. For all remaining accessions, any heterozygous genotype call was transformed to missing data, and accessions with > 1% missing data among all SNPs were excluded.

### Patterns of chloroplast and chromosome 1 translocation polymorphism

For chloroplast, bi-allelic SNPs of 67 protein coding genes were called using vcftools^25^ after conversion of fasta alignment format to vcf format in TASSEL^26^. Focusing on bi-allelic sites with no missing data and zero heterozygosity, we identified 760 sites polymorphic within *A. thaliana* and 2124 fixed between *A. thaliana* and any one outgroup species. SNPs were identified in chromosome 1 translocation following procedures of previous studies^2,16^. PCA was done with R package adegenet^27^ and visualized with R package ggplot2^28^.

Accessions of *Arabidopsis thaliana* were assorted into groups (geo-clusters) according to geographical location where they were sampled (Supplementary Table 3). Diversity of each chloroplast within each geo-cluster was then estimated by pair-wise genetic distance^29^. To account for uneven sampling, the average genetic variation of 100 resampling trials was obtained for each geo-cluster. For each re-sampling trial, 100 samples were randomly drawn with replacement within each geo-cluster. Groups with less than 3 samples in each geo cluster were ignored. All the calculation and plots were completed in R with customize scripts and package ggplot2^28^. Pie charts were plotted with R package ggplot2^28^ as well.

### Phylogenetic reconstruction and divergence time estimation

5,915,870 nuclear SNPs of 1323 accessions (including *Arabidopsis lyrata*) were used to construct nuclear neighbor-joining tree. The pair-wise distance is calculated by dividing number of SNP difference between pairs with total number of non-missing sites that are polymorphic within 1323 accessions.

1312 chloroplast haplotypes of 2124 bi-allelic SNPs were converted into phylip format and submitted to maximum-likelihood-based phyML 3.0^30^ for reconstruction of phylogenetic tree. Substitution model selection was done by SMS^31^ with Bayesian Information Criterion (BIC). Subtree pruning regrafting (SPR) was used as tree searching algorithm and the branch support was estimated by approximate likelihood ratio test (aLRT SH-like). Branches with low support (aLRT < 0.5) were collapsed with TreeGraph2^32^. The collapsed maximum likelihood tree was visualized and colored in FigTree version 1.4.3 (http://tree.bio.ed.ac.uk/software/figtree/).

Divergence time among haplogroups was estimated with BEAST version 2.5.0^12^. The collapsed ML tree of 426 unique chloroplast haplotypes was constructed as described previously and served as starting tree for BEAST after it was converted ultrametric and had the node ages fit within constraints of calibration points using R package ape^33^. Three calibration points estimated in previous studies^7,11^ were adopted, the root height (divergence time between genus *Arabidopsis* and *Camelina, Capsella*) was set to 8.16 Mya; divergence time between genus *Capsella* and *Camelina* was set to 7.36 Mya; and the divergence time between *Arabidopsis thaliana* and other species in *Arabidopsis* genus was set to 5.97 Mya. Normal distribution with 1 mya standard deviation was used for all the three calibration points.

Two independent MCMC runs with 5 x 10^8^ chain length were generated with Calibrated Yule Model for priors and Relaxed Clock Log Normal for clock model. GTR was chosen as site model according to SMS. Parameters of MCMC trees were sampled every 5 x 10^4^ generations and submitted to Tracer version 1.6 (http://tree.bio.ed.ac.uk/software/tracer/) for quality control of MCMC chains. LogCombiner^12^ was implemented to combine two independent runs. A maximum clade credibility (MCC) tree was constructed using 18000 output trees of LogCombiner with 10% burn-in in TreeAnnotator^12^. The MCC tree was then visualized and colored in FigTree version 1.4.3 (http://tree.bio.ed.ac.uk/software/figtree/).

### ABBA-BABA and 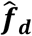 estimation

Python and R scripts were downloaded from (https://github.com/simonhmartin/genomics_general) for ABBA-BABA and 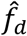 estimation^17^. Genome-wide D statistics were estimated in R followed the instruction of (http://evomics.org/learning/population-and-speciation-genomics/2018-population-and-speciation-genomics/abba-baba-statistics/), West European population (EU) was treated as Pop1, Yangtze population (YA) was treated as Pop2, Yunnan (YU) (the less admixed accession) was treated as Pop3 and *Arabidopsis lyrata* was treated as outgroup. Sliding window analysis of 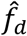 estimation between Yangtze population and Yunnan (the less admixed accession) was done using the Python scripts. Window size was set to 50 kb with 20 kb step, each window containing at least 100 SNPs. Windows with top 5 % highest 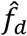 values were viewed as regions with strong introgression between Yangtze and Yunnan. The selected genes in Yangtze population^8^ within/outside the window of introgression were then submitted to agriGO v2.0^34^ for gene ontology analysis.

### Multiple sequentially Markovian coalescent analysis (MSMC)

Relative cross coalescence rate was estimated using MSMC v2^15^. Sequences of chromosome 1 translocation (Chr1:20271447-21032307) were used as input. Generation time was set at 1 year and mutation rate of 7.1 x 10^-9^ was assumed according to previous studies^5,8^. Results were then plotted in R.

### Extended Bayesian skyline plot

Extended Bayesian Skyline analysis was done using BEAST v2.5.0^12^ with fixed number of 2124 chloroplast genome polymorphisms. Parameters were set in BEAUti 2^12^, HYK substitution model with empirical frequency was chosen and the clock model was set to strict clock with clock rate estimated by previous Calibrated Yule Model (0.0223). Priors were set to Coalescent Extended Bayesian Skyline with 0.5 population model factor and default value for the rest of parameters. Sufficient length of MCMC chains were run to achieve acceptable ESS values, which indicates the model is well-mixed. The ESS values were estimated in Tracer v1.6.0.^12^, and the results were plotted in R.

## Acknowledgements

We thank Thomas Mitchell-Olds and the Lee lab member for comments to this manuscript. We thank all researchers who have made the *Arabidopsis* genetic resources and data publicly available. We are grateful to Computer and Information Networking Center, National Taiwan University for the support of high-performance computing facilities. This work is supported by the Ministry of Science and Technology of Taiwan (105-2311-B-002-040-MY2 and 107-2636-B-002-004 to CRL).

## Author contributions

Designed the study: CRL. Analyzed data: CWH, CRL, CYL. Wrote the paper: CRL, CWH.

## Conflict of interest statement

The authors declare no conflict of interest

## Data Accessibility

All data were downloaded from public database.

---

**Supplementary Fig. 1.**
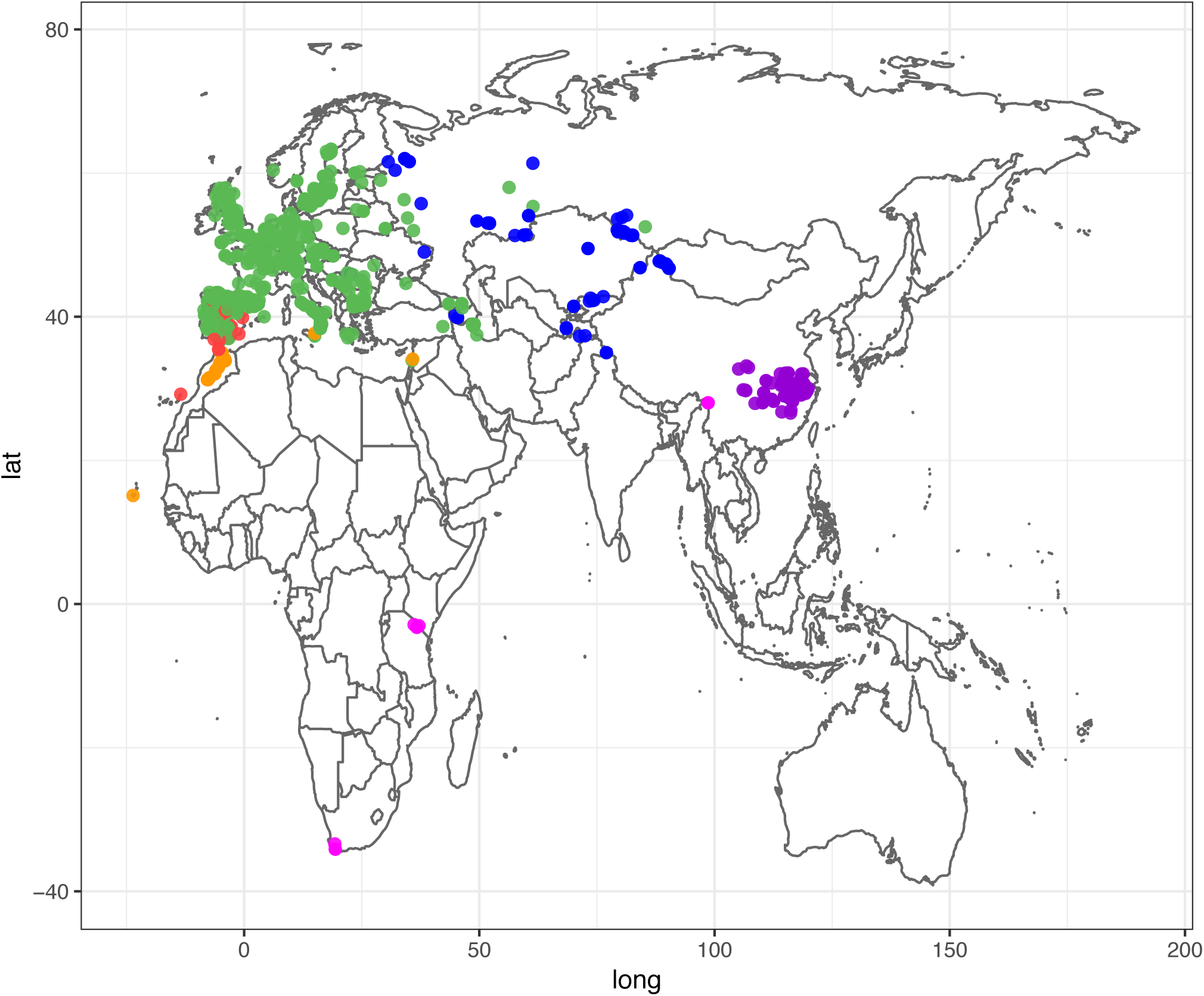
Geographical distribution of nuclear genetic variation in *Arabidopsis thaliana*. The color of dots corresponds to the Neighbor-Joining tree in Figure 1a.

**Supplementary Fig. 2.**
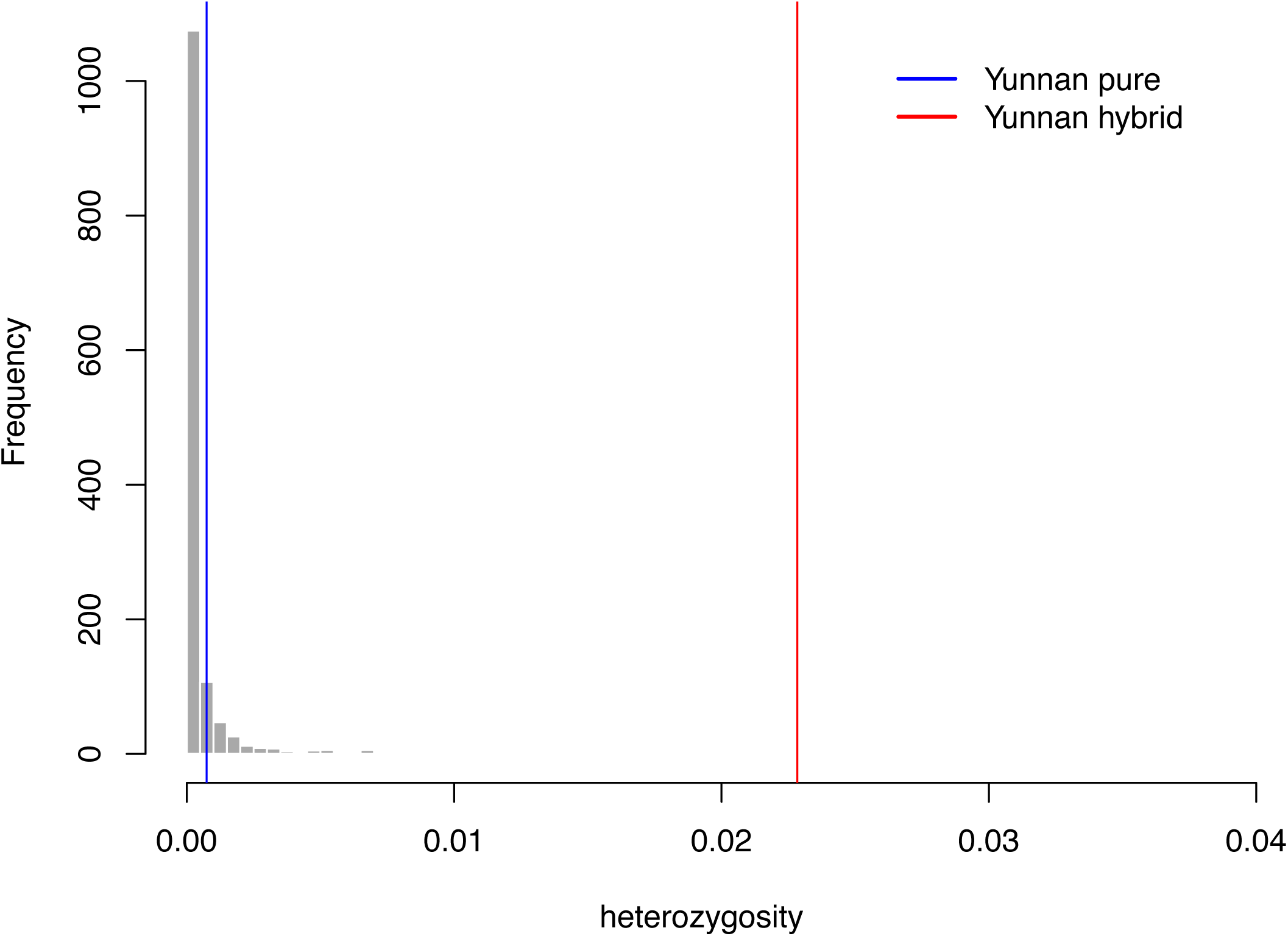
Distribution of nuclear genome heterozygosity of 1322 *Arabidopsis thaliana* accessions.

**Supplementary Fig. 3.**
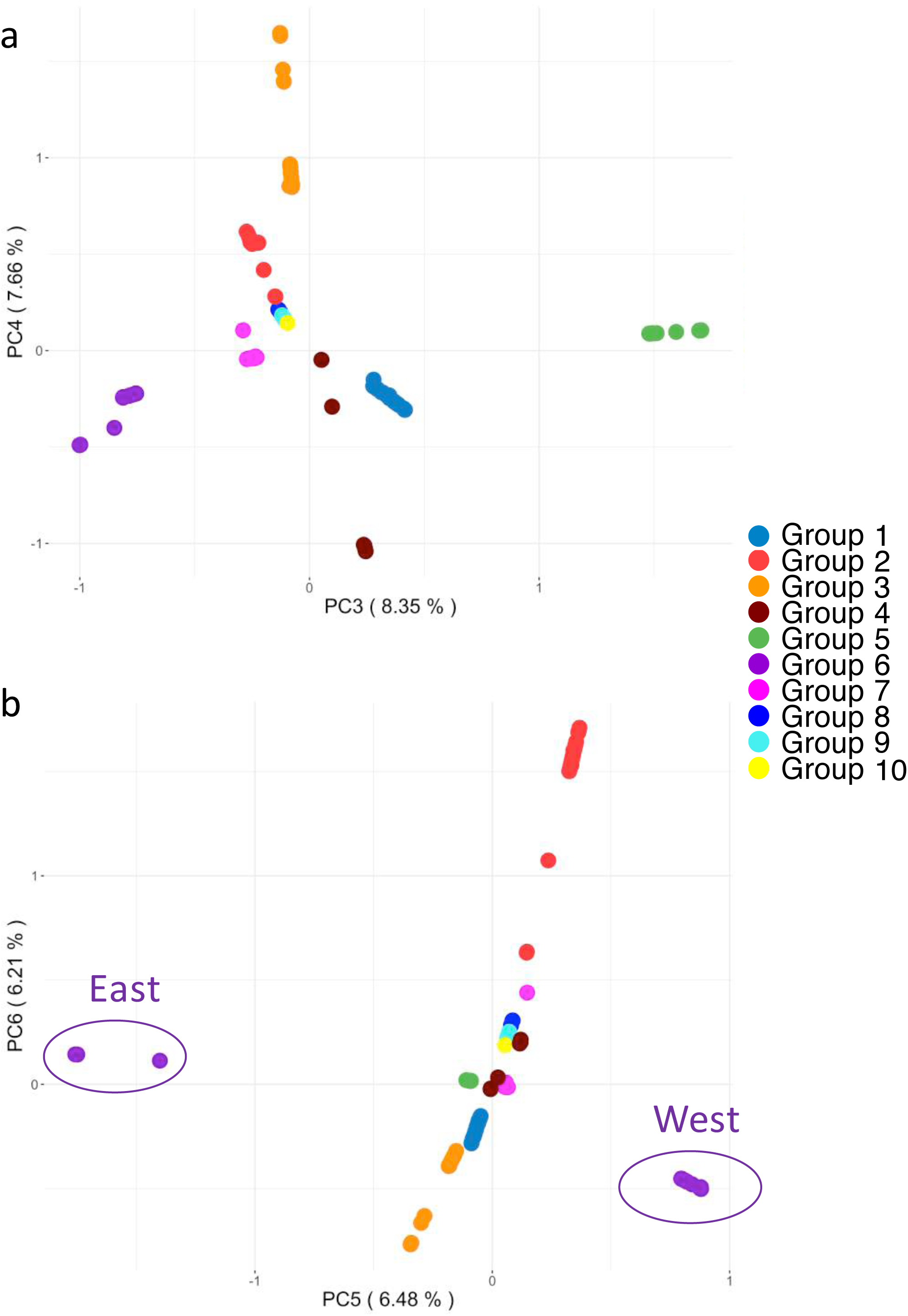
Differentiation of *Arabidopsis thaliana* chloroplast genomes. (a) PC3 and PC4. (b) PC5 and PC6. Group 6 (East) consists of samples from Yangtze River Basin, China and Kashmir, India. Group 6 (West) consists of samples in Eurasia.

**Supplementary Fig. 4.**
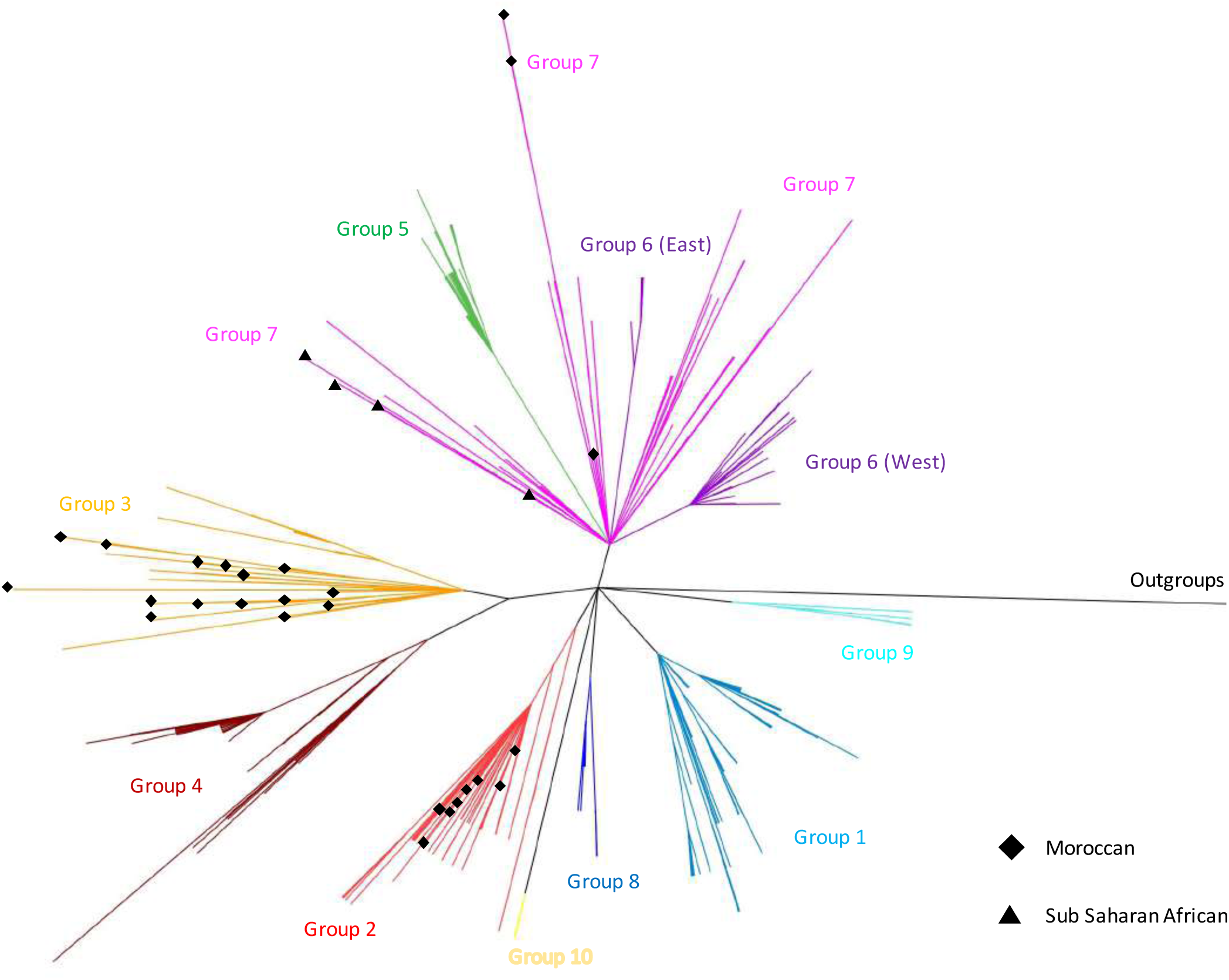
Chloroplast uncollapsed maximum likelihood phylogram. Group 6 (East) consists of samples from Yangtze River Basin, China and Kashmir, India. Group 6 (West) consists of samples in Eurasia. Note that this is an uncollapsed bifurcating tree. Some internal branches are too short to be clearly visible. These branches also tend to have extremely low branch support.

**Supplementary Fig. 5.**
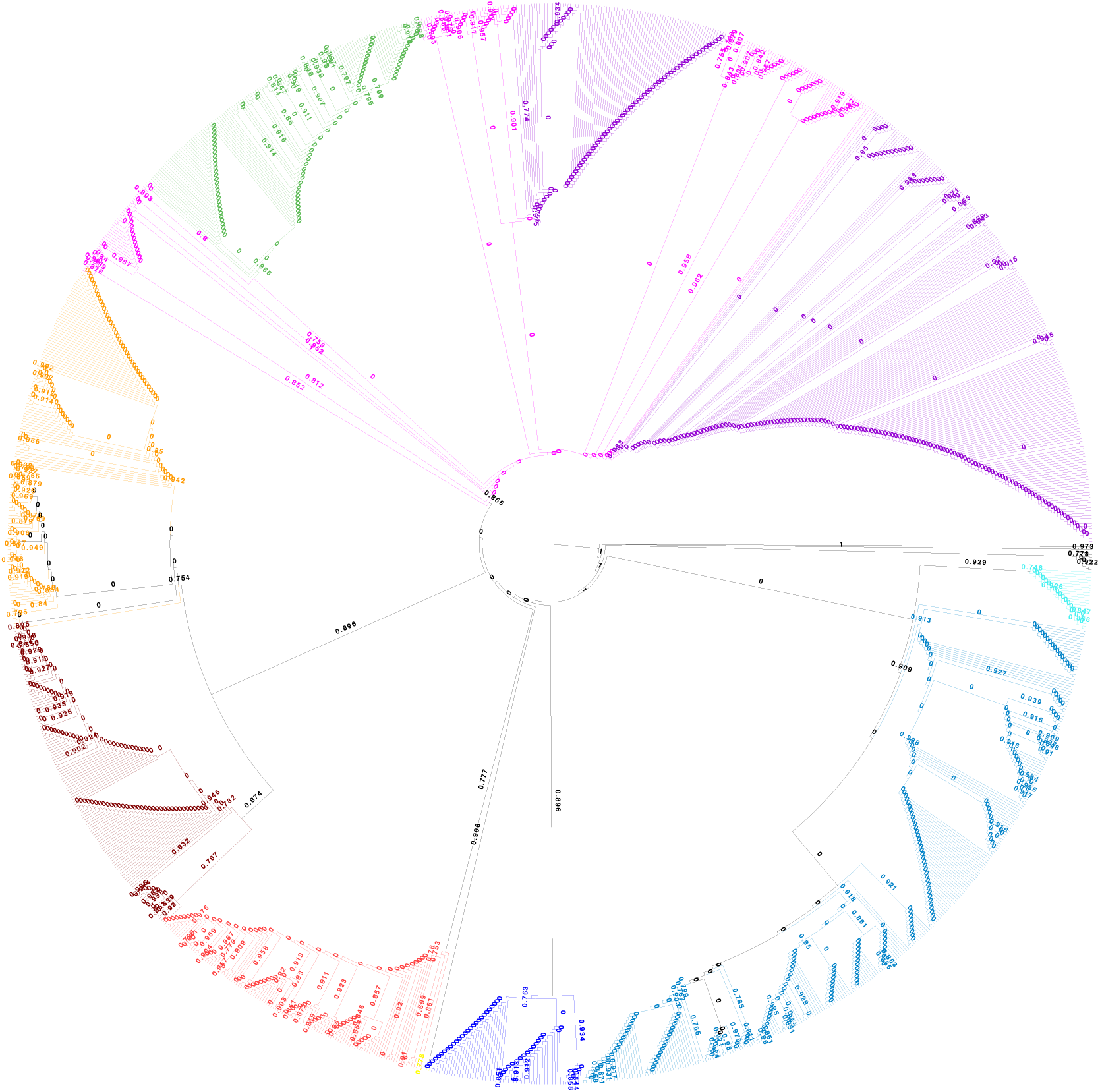
Chloroplast uncollapsed maximum likelihood cladogram with aLRT branch support. Branches were colored according to chloroplast haplogroups defined in Figure 2.

**Supplementary Fig. 6.**
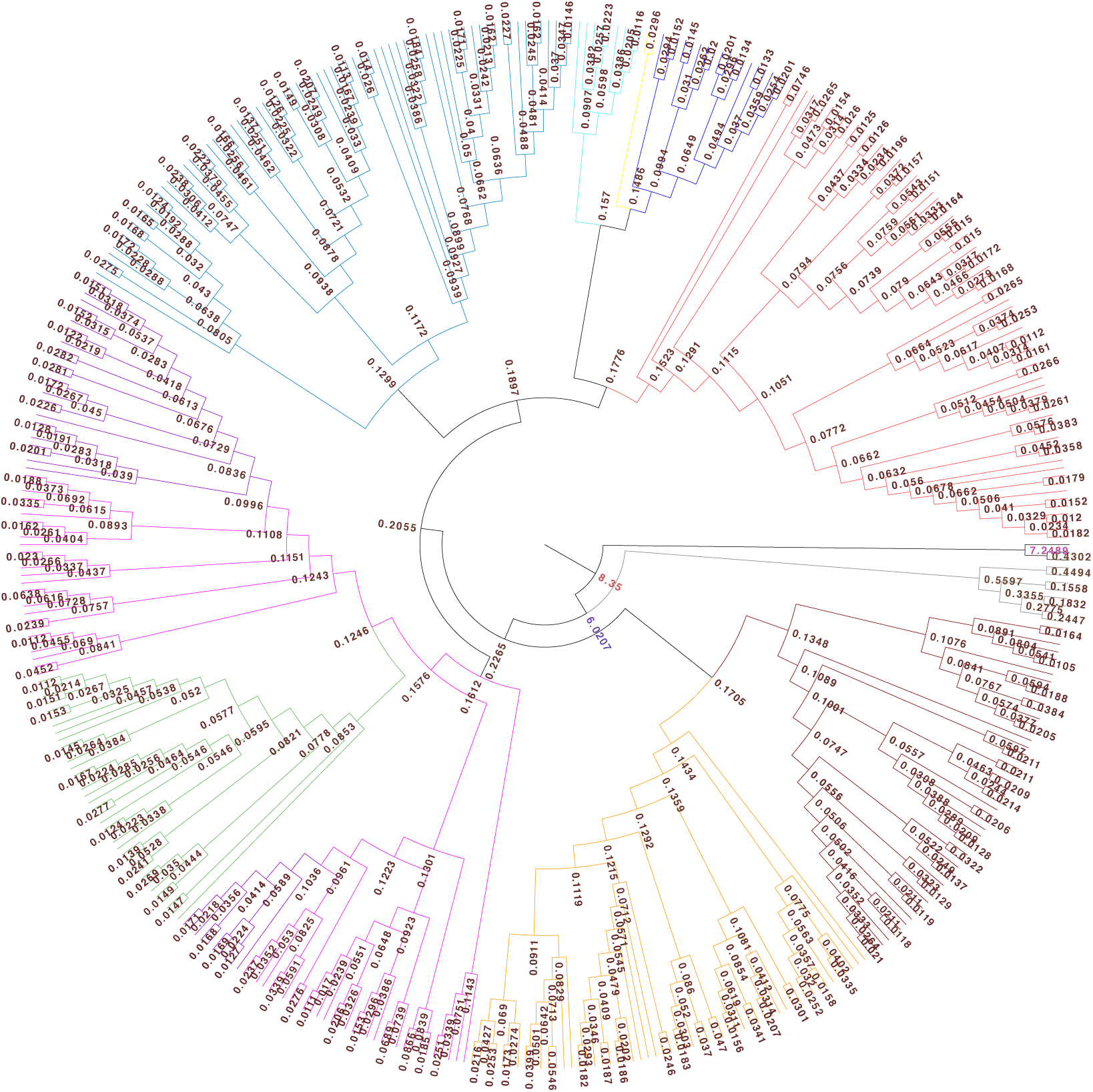
Chloroplast BEAST dated tree. Node values represent mean height of divergence time in mya. Branches were colored according to chloroplast haplogroups defined previously.

**Supplementary Fig. 7.**
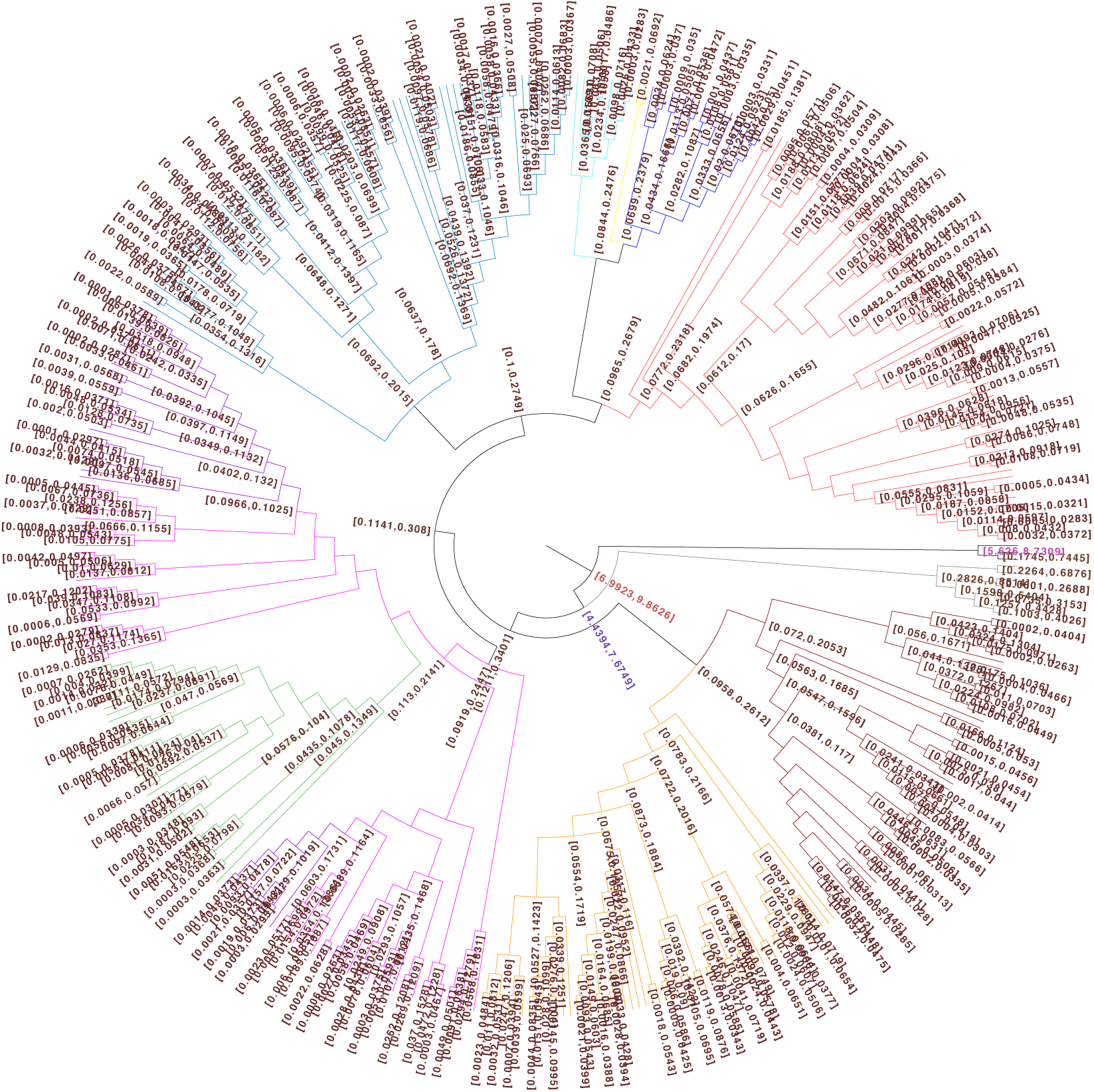
Chloroplast BEAST dated tree. Node values represent 95% highest posterior density range of divergence time in mya. Branches were colored according to chloroplast haplogroups defined previously.

**Supplementary Fig. 8.**
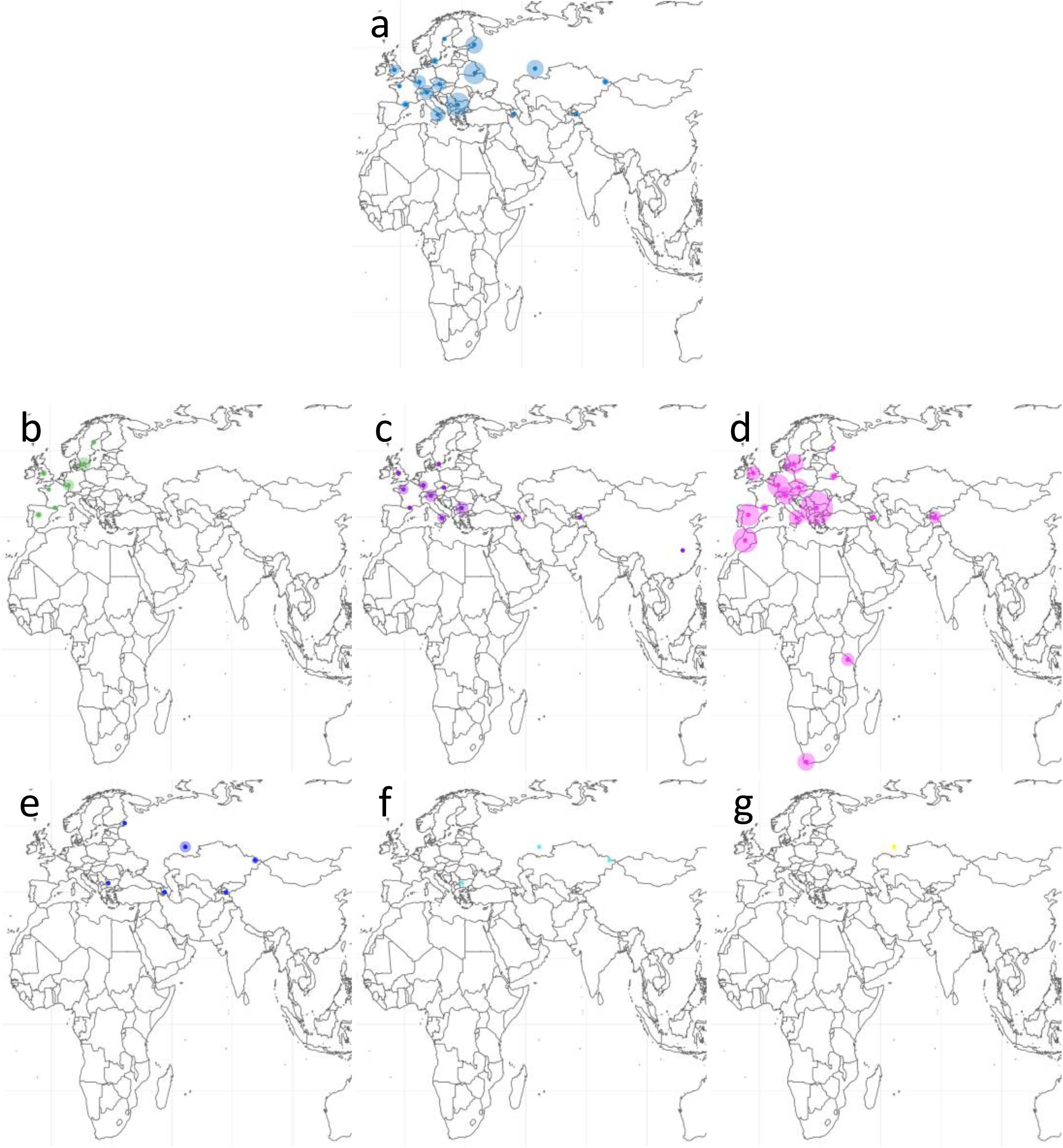
Spatial genetic variation of chloroplast haplogroups. (a), (b), (c), (d), (e), (f), (g) are polymorphism maps correspond to group 1, 5, 6, 7, 8, 9, 10 respectively. The diameter of each circles is proportional to mean pair-wise genetic distance of each geographical region.

**Supplementary Fig. 9.**
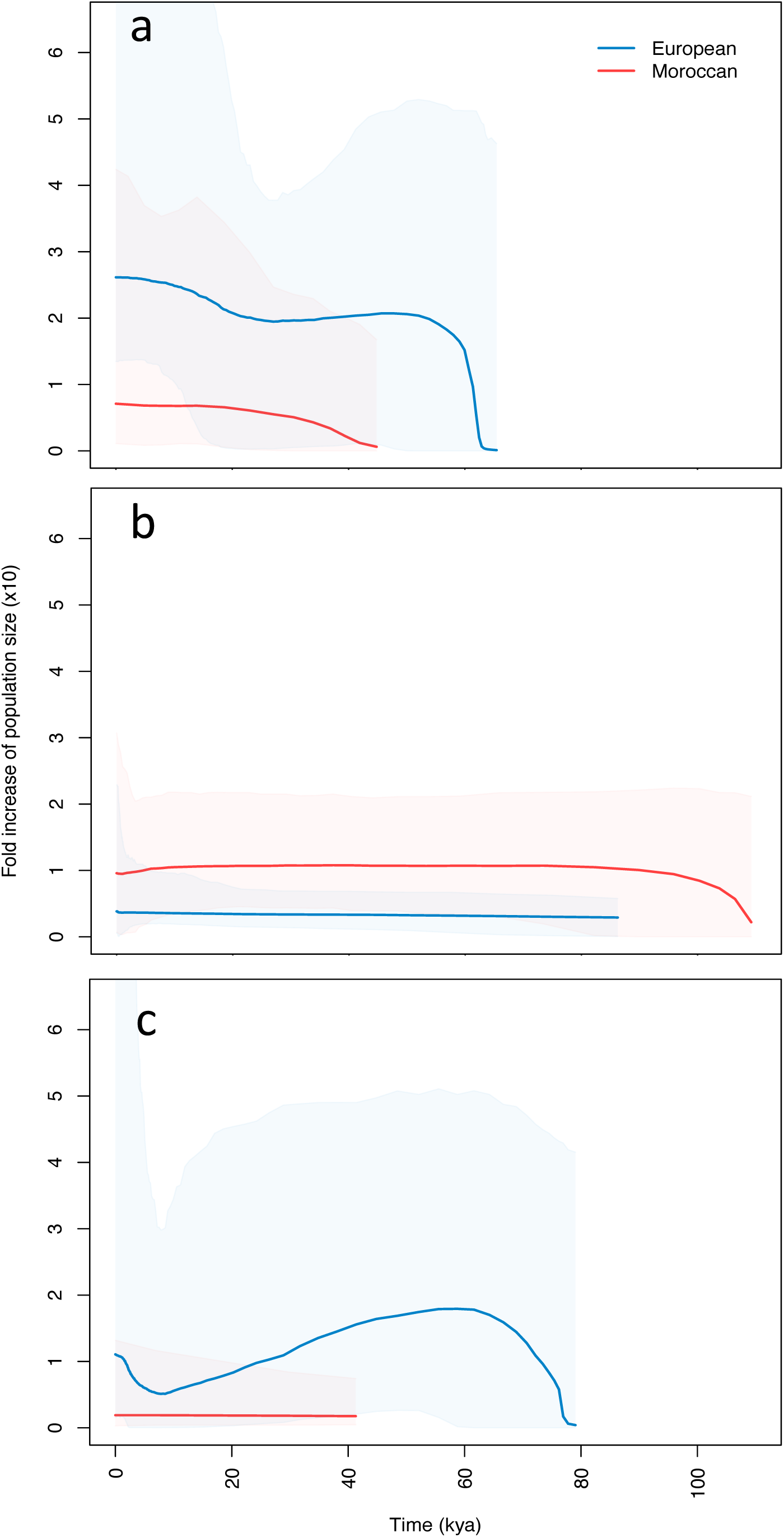
Variation of population size over time inferred from chloroplast polymorphism. Extended Bayesian Skyline is plotted for European and Moroccan population of chloroplast (a) haplogroup 2, (b) haplogroup 3 and (c) haplogroup 7.

**Supplementary Fig. 10.**
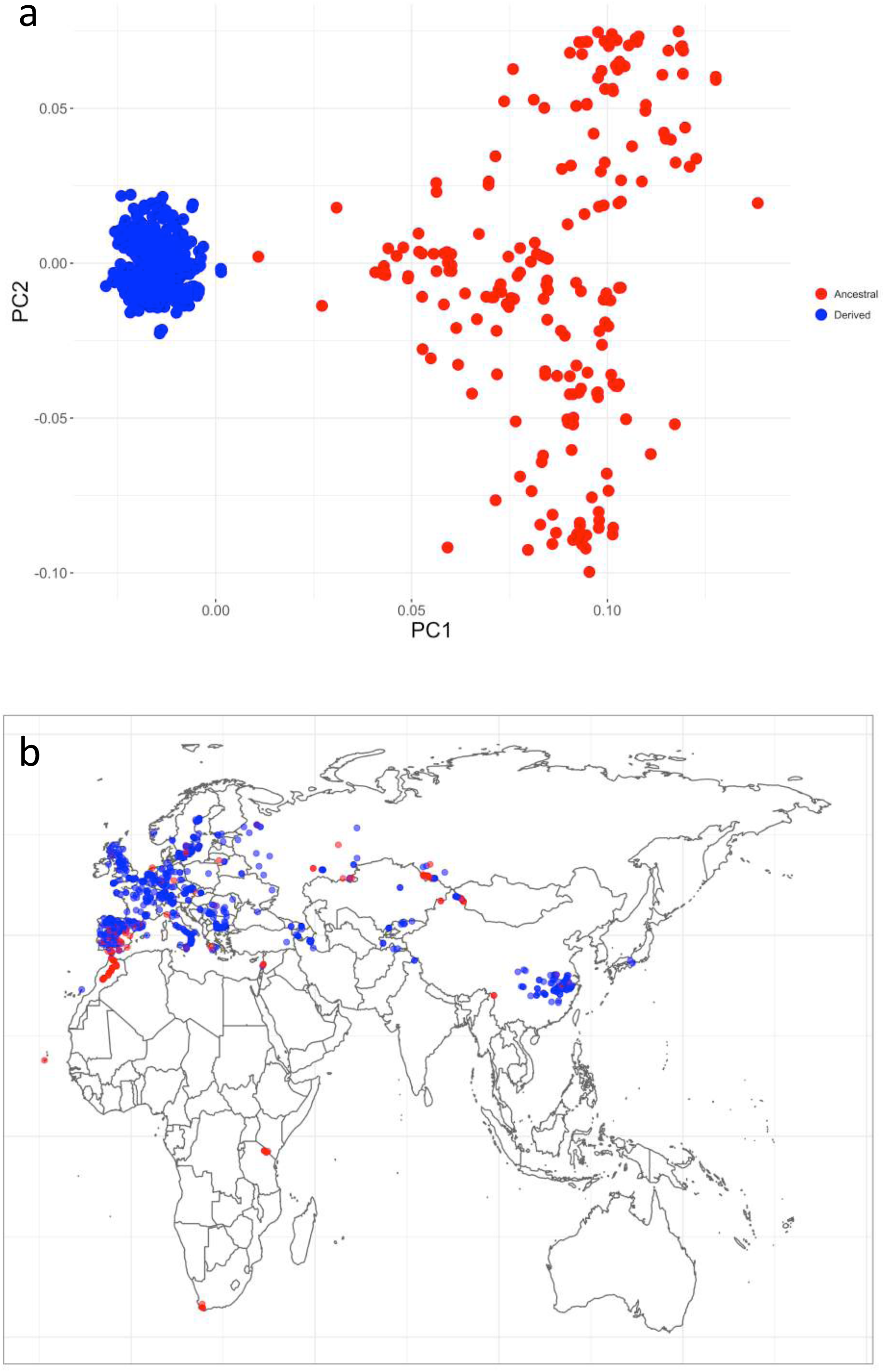
Genetic differentiation of chromosome 1 translocation. (a) Principal component analysis. (b) Geographical distribution. Red dots are accessions with the ancestral haplotype, and blue are accessions with the rearranged derived haplotype.

**Supplementary Fig. 11.**
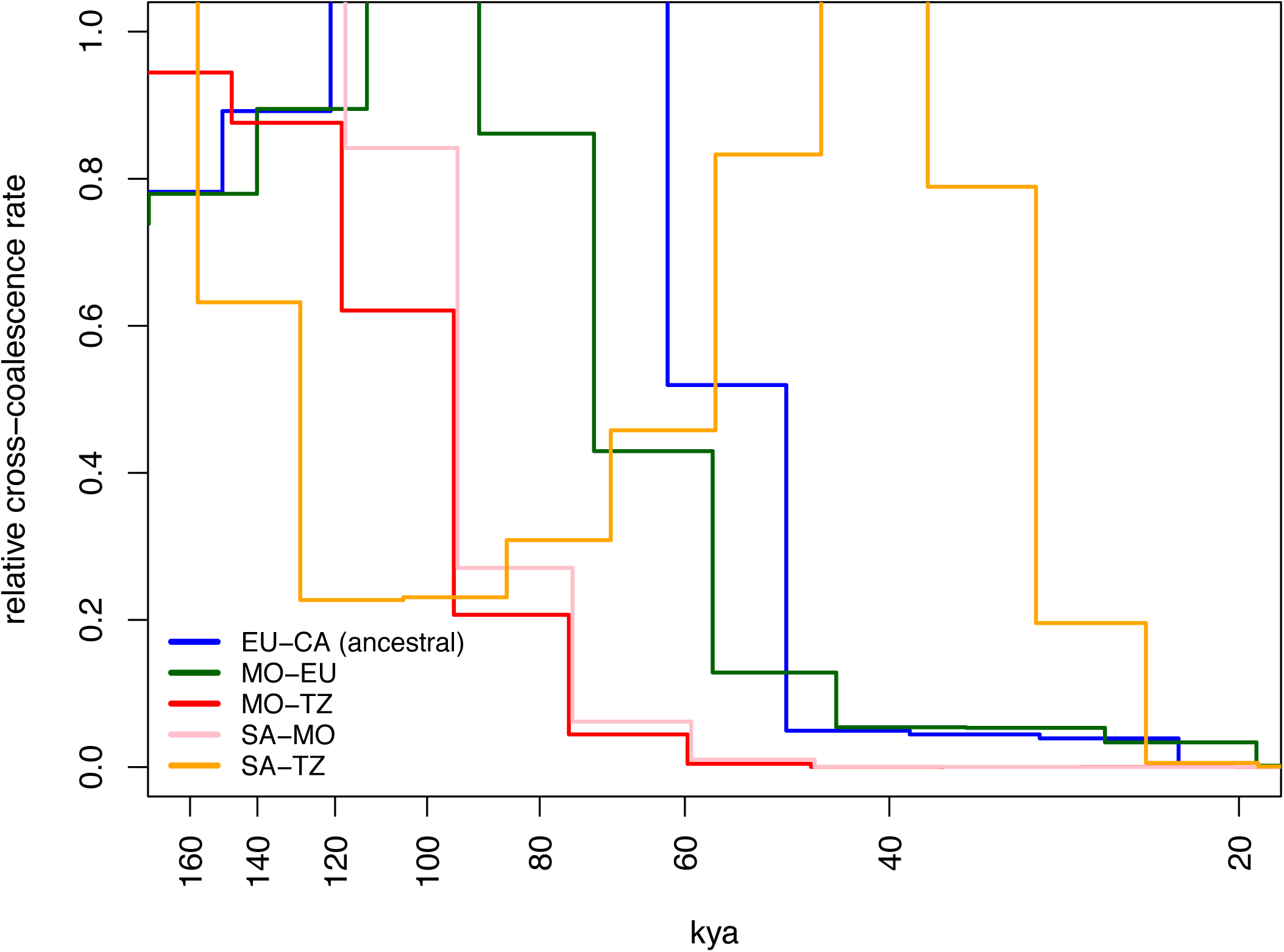
Reproducing 4-haplotype MSMC results of Durvasula *et al*. (2017) using chromosome 1 translocation instead of whole genome. Relative cross coalescence rate (CCR) between populations is shown: EU: West Europe, CA (ancestral): Central Asia accession with ancestral allele of chromosome 1 translocation, MO: Morocco, TZ: Tanzania, SA: South Africa. Decrease of CCR from 1.0 indicates population split, steep slope of CCR from 1.0 to 0.0 indicates drastic and complete isolation while mild one indicates slow and progressive isolation.

**Supplementary Fig. 12.**
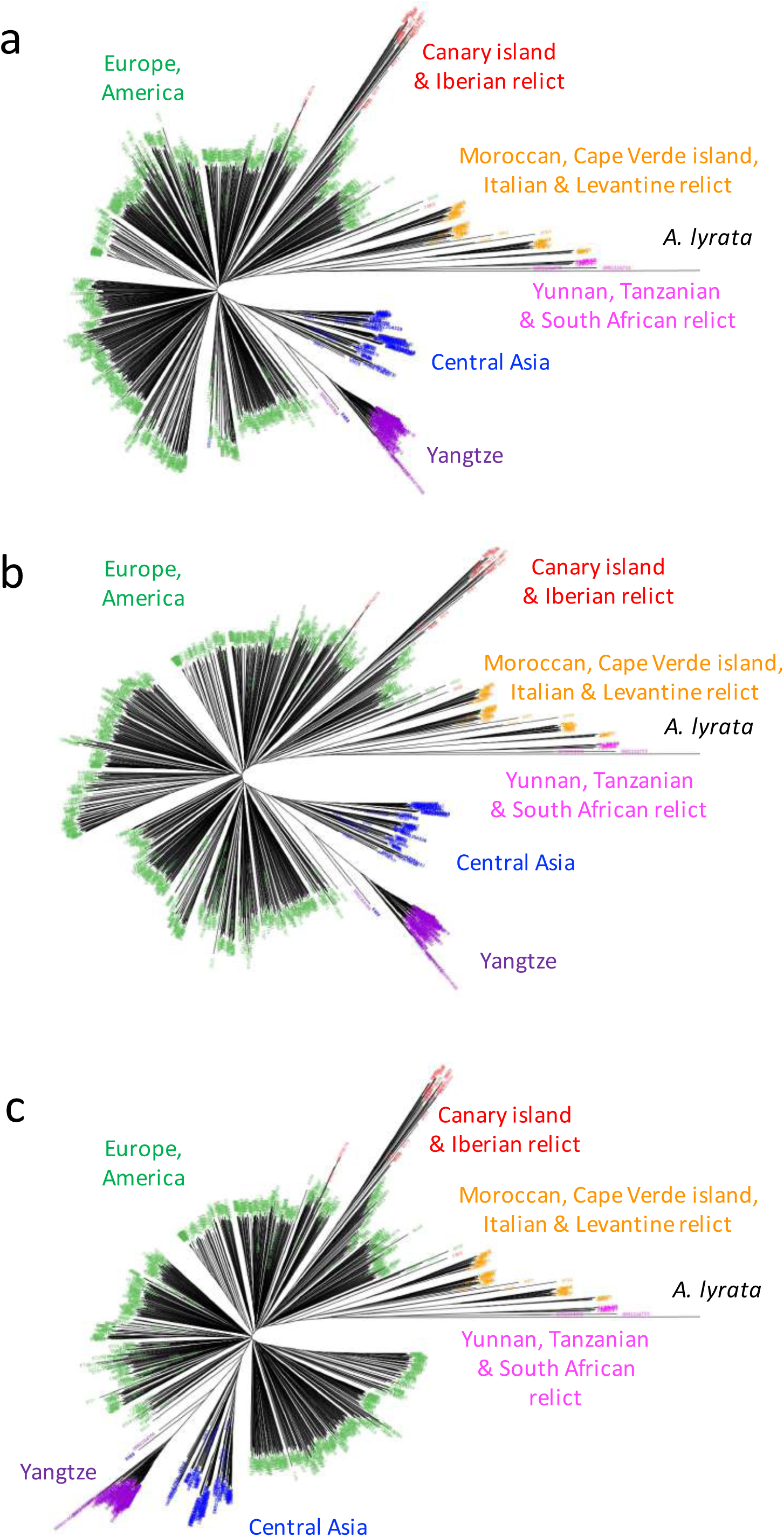
Nuclear Neighbor-Joining tree built (a) without SNPs located in regions of top 20% 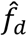, (b) without SNPs located in regions containing selected genes of Yangtze population and (c) both.

